# Design principles for accurate folding of DNA origami

**DOI:** 10.1101/2024.03.18.585609

**Authors:** Tural Aksel, Erik J. Navarro, Nicholas Fong, Shawn M. Douglas

## Abstract

We describe design principles for accurate folding of three-dimensional DNA origami. To evaluate design rules, we reduced the problem of DNA strand routing to the known problem of shortest-path finding in a weighted graph. To score candidate DNA strand routes we used a thermodynamic model that accounts for enthalpic and entropic contributions of initial binding, hybridization, and DNA loop closure. We encoded and analyzed new and previously reported design heuristics. Using design principles emerging from this analysis, we redesigned and fabricated multiple shapes and compared their folding accuracy using electrophoretic mobility analysis and electron microscopy imaging. We demonstrate accurate folding can be achieved by optimizing staple routes using our model, and provide a computational framework for applying our methodology to any design.

---

In 2006, Rothemund pioneered the DNA origami technique, which utilizes a single-stranded DNA (ssDNA) ‘scaffold’ and numerous short DNA oligonucleotide ‘staples’ to orchestrate the self-assembly of predetermined shapes^1^. This versatile approach to bottom-up fabrication^2–4^ holds the potential to revolutionize manufacturing of nanoscale materials and devices^5–8^, including tiny mechanical instruments for biological and biophysical measurements^9–13^, and programmable therapeutics^14–16^. A bottleneck to realizing many applications with current design approaches is folding accuracy, or how closely the structure of a folded DNA origami shape agrees with the structure of the intended design^17^.

Determining generalizable design principles that increase the folding accuracy of DNA origami has been recognized as an important and unsolved challenge^18–23^. With the goal of unlocking high-precision applications of the method, we developed a computational framework that addresses three related problems: how to express design principles as rules that are both human- and machine-readable, how to score the quality of a candidate staple route, and how to implement design rules in a fast, global, and unbiased fashion. We applied our framework to explore several aspects of DNA strand design, and report design principles emerging from this analysis.

Our approach maps an input design onto a weighted graph in which smaller edge weights correspond to better-scoring staple routes, and therefore a *shortest-path* algorithm can be applied to find a globally optimized set of staple routes. We considered several schemes for scoring the quality of a staple route and observed favorable results using a thermodynamic model that accounts for staple-to-scaffold binding, scaffold loop closure, and hybridization. We used our algorithm to refine several complex lattice-based structures and noted enhanced folding accuracy across all designs when assessed by gel electrophoresis and transmission electron microscopy (TEM). We encapsulated our design principles and algorithm in a computational notebook to enable researchers to analyze and optimize any design.

## Reduction to a known problem

A DNA origami design can be understood as a directed graph in which each node represents a nucleotide and edges represent phosphate-backbone or base-pairing connections between nucleotides. Cadnano, a computer-aided design tool, provides a graphical interface to enable users to manipulate a structured layout of this directed graph^24^. Although the directed graph layout provides a useful molecular abstraction, the format is ill-suited for analyzing the underlying design principles. Therefore, we mapped the directed graph onto a separate weighted graph in which nodes represent staple precursor phosphates. Edges in the second graph represent possible design choices that would remove the phosphates at the connected nodes. Edge weights are assigned by a scoring function that estimates the quality of the staple flanked by the removed phosphates. In this framework, paths are sets of design choices that can be mapped back to the directed graph to generate staple paths, and the shortest paths represent globally optimized solutions according to the scoring function.

Our semi-automated approach consists of creating an incomplete design in Cadnano, and then using a computer program to carry out the mapping, staple-route selection, and reverse-mapping steps to generate an output. In the manual step, users draw a scaffold route and invoke the “autostaple” function to generate precursor staples—long, often circular, staple routes complementary to the scaffold. Precursor staples can be customized as needed before saving the design as a Cadnano JSON file.

The automated step begins by parsing the input file to construct a directed graph that captures key details of the design: scaffold and staple routes, base-pairing relationships, and the locations of strand termini, strand crossovers, base insertions, and base skips (Fig. 1A). Next, staple precursors found in the directed graph are mapped onto a separate weighted graph whose nodes represent potential breakpoints, or phosphate bonds between nucleotides. Thus, each edge represents a candidate staple route, connecting pairs of breakpoint nodes that, if broken, would yield a valid staple. Each staple precursor forms a subgraph within the weighted graph.

**Figure 1.**
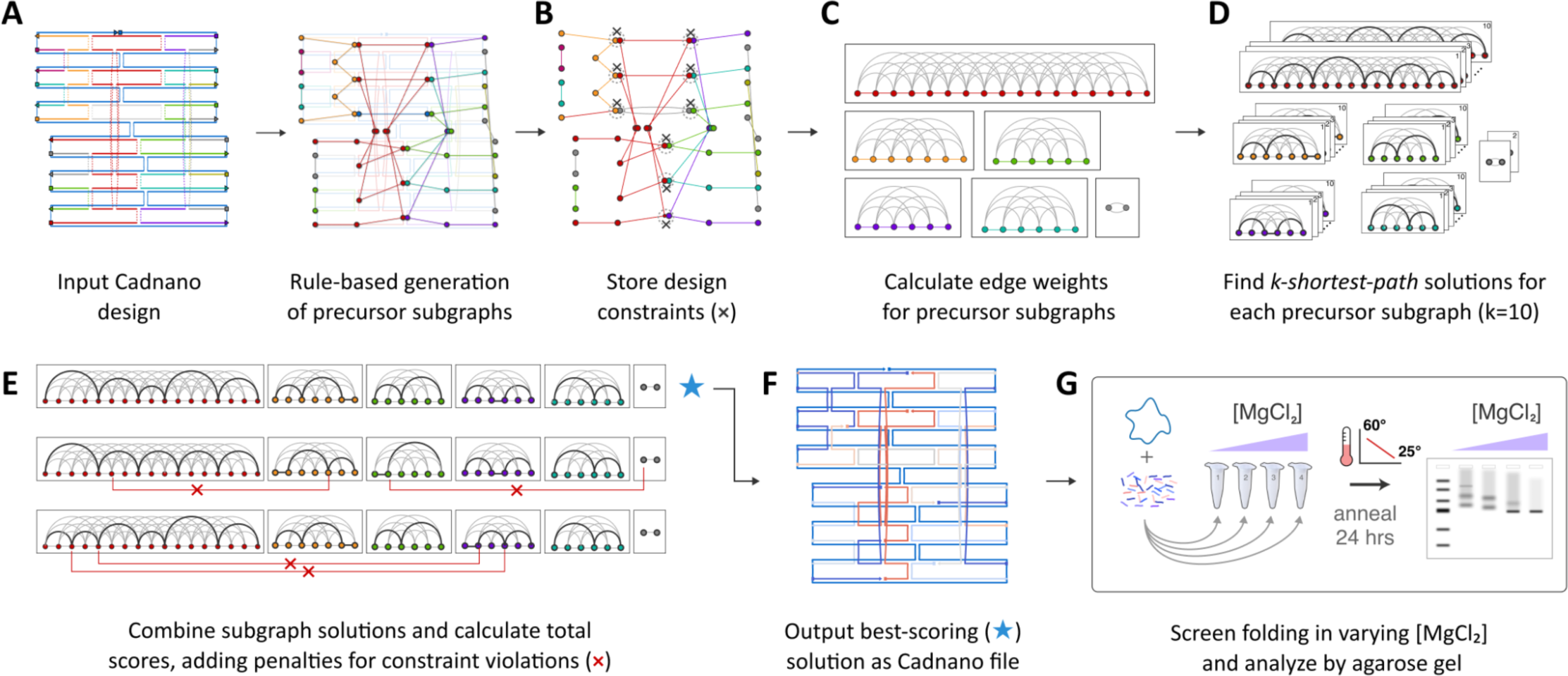
Overview of our staple-routing algorithm. (**A**) The user prepares a Cadnano design, leaving auto-generated staple precursors unbroken. Based on user input for allowed staple breakpoints, our script maps the input design to a weighted graph in which nodes represent candidate breakpoints, and edges represent staples that terminate at the flanking breakpoints. Each staple precursor maps to an individual subgraph. (**B**) Graph constraints are stored in order to later penalize route solutions that violate design intent, e.g., break both halves of a crossover. (**C**) Edge weights are calculated using the designated scoring function. (**D**) The *k-shortest-paths* algorithm is applied to the precursor subgraphs. (**E**) Subgraphs are assembled into candidate solutions, and constraint violations are penalized. (**F**) The best-scoring solution is used to output a Cadnano file implementing the corresponding staple routes. The output staple colors correspond to normalized values derived from the scoring function. (**G**) Design performance is analyzed using a folding performance screen followed by agarose gel electrophoresis.

The weighted graph is pruned during construction using a rule-based approach to include or exclude specific types of nodes or edges, thereby implementing design rules in a globally consistent manner. The breakpoint rules we explored are shown in Fig. S1. Edges are pruned to ensure final staple lengths are 21 to 60 nt by default. Design constraints are introduced into the weighted graph to later calculate penalties for global solutions that violate the original design intent by breaking both halves of any double crossover (Fig. 1B).

Next, a scoring function is used to assign edge weights (Fig. 1C). Our modular approach can use any function that takes a staple route input and returns a numerical score. Our preferred scoring function is described below. The weighted subgraphs are analyzed using a *k-shortest-path* algorithm to identify the *k* shortest path solutions for each subgraph, with *k=10* by default (Fig. 1D). Finally, a set of candidate global solutions is assembled. Each global solution is constructed by selecting, at random, one of the *k* solutions for each subgraph (Fig. 1E). Penalties are assessed for subgraph combinations that violate design constraints. The total score for a global solution is calculated as the sum of edge weights for all staples in the design. The best-scoring global solution is reverse-mapped to the directed graph to generate a final Cadnano design, which is output to disk along with a detailed score report (Fig. 1F). Designs can then be synthesized, folded, and subjected to experimental analysis (Fig. 1G).

### Designing a scoring function

Shortest-path optimization compares the relative scores of candidate staple-routing solutions and returns the best-scoring result. However, better scores are only meaningful if they correlate with improved folding accuracy. We aimed to devise a scoring function that encapsulated design parameters that affect the probability of successful folding. We adopted an equilibrium modeling approach by independently calculating each staple score in its fully-bound state, and ignoring folding kinetics and inter-staple cooperativity. The probability of an origami design folding accurately, *P_origami_*, is the product of the probabilities of folding of each staple *P_staple_* in the design, where *n* is the total number of staples in the design.

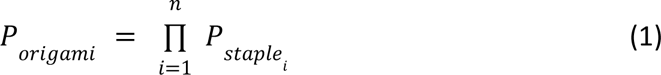

We used a Boltzmann distribution to express *P_staple_* in terms of *ΔG_total_*, the Gibbs free-energy change between the unfolded and folded states, with *R* and *T* denoting the molar gas constant and temperature, respectively:

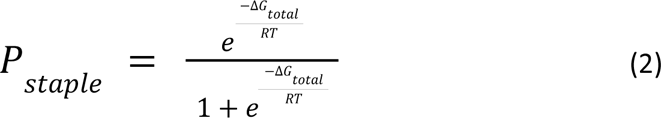

We assigned edge weights as logarithmic values of *P_staple_*. A global quality score, *Q_origami_*, is the sum of the edge weights normalized by *L*, the total number of base pairs formed by scaffold-staple duplexes:

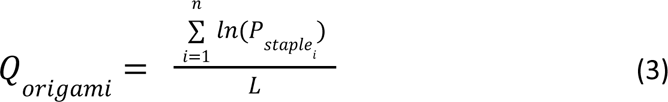

As summarized in Fig. 2B, we estimated *ΔG_total_* for each candidate staple route as the sum of three energy terms:

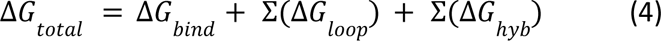

**Figure 2.**
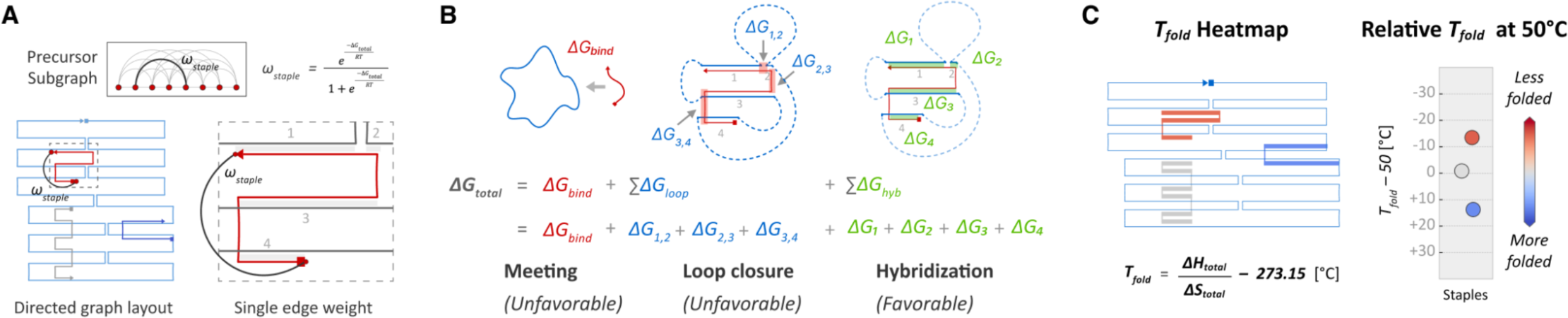
A three-term thermodynamic model is used to calculate edge weights. (**A**) In the precursor subgraphs, edges connect pairs of breakpoint nodes (red circles) that flank a possible staple route. Here, candidate staple (boxed) is complementary to four distinct stretches of the scaffold (1, 2, 3, 4). (**B**) *ΔG_total_*values are calculated from the sum of three thermodynamic terms representing the meeting of the staple and scaffold (*ΔG_bind_*), each “loop closure” that constrains the 3D movement of two non-contiguous stretches of the scaffold (*ΔG_loop_*), and the free-energy gain due to hybridization between the staple and scaffold (*ΔG_hyb_*). (**C**) Optimized solutions are reversed-mapped to a ‘heatmap’ directed-graph layout with staples colored by the predicted folding temperature *T_fold_*. A strip plot displays the *T_fold_*values relative to 50°C. Staples predicted to be less folded at T=50° are colored red; more-folded staples are colored blue.

The *ΔG_bind_*term is the concentration-dependent free-energy change due to the bimolecular binding event when the staple first associates with the scaffold ^19^. The second term, *ΔG_loop_*, is the energy cost for each scaffold loop closure whereby the staple restricts movement of non-contiguous scaffold segments. Like Dunn *et al*. ^19^, we treat the scaffold as a freely jointed chain, and calculate the mean-squared distance between the two scaffold segments to be bridged by the staple. The distance is used to estimate the effective local concentration of one end of the loop at the other and then calculate the *ΔS_loop_*and *ΔG_loop_*values. The third term, *ΔG_hyb_*, is the free-energy gain due to hybridization of the staple to the scaffold ^25,26^. Free energy calculations were performed at T=50°C, where nearest-neighbor predictions of ΔG° are most accurate ^25^, and near the temperature intervals where DNA origami folding has been observed empirically ^27^. We calculated enthalpic (*ΔH_total_*) and entropic (*ΔS_total_*) components for each staple and used the Gibbs free-energy equation to estimate a folding temperature, *T_fold_,* for each staple when *ΔG_total_* = 0.

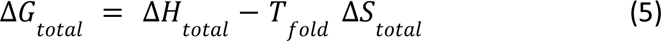

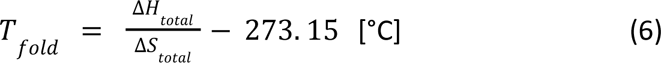

We used *T_fold_* estimates for each staple to generate a ‘heatmap’ representation using the conventional directed-graph schematic (Fig. 2C). Staples are assigned colors using a cool-warm colormap, with a center point at 50°C. Values of *T_fold_* > 50°C appear blue, *T_fold_* ≈ 50°C are gray, and *T_fold_* < 50°C appear red.

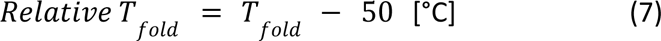

We visualized relative *T_fold_* values by subtracting 50°C from the absolute *T_fold_*, as shown in Eqn 7.

### Validation

To validate our approach, we redesigned four published multilayer DNA origami shapes, and assessed their folding accuracy compared to the original designs using agarose gel electrophoresis and TEM (Fig. 3). The panel of objects included a 100-helix block (10×10 layout) with square-lattice packing, and three different 64-helix blocks (4×16, 8×8, and 16×4 layouts) each with honeycomb-lattice packing^18,24^. We added a ‘read-only’ mode to our algorithm that applies the scoring function to existing designs without breaking any staples and used it to generate *T_fold_*heatmaps (Fig. 3 col. 2, Figs. S2–S5) and strip plots (Fig. 3 col. 3). The estimated mean *T_fold_*values for the original were uniformly lower than the redesigned versions.

**Figure 3.**
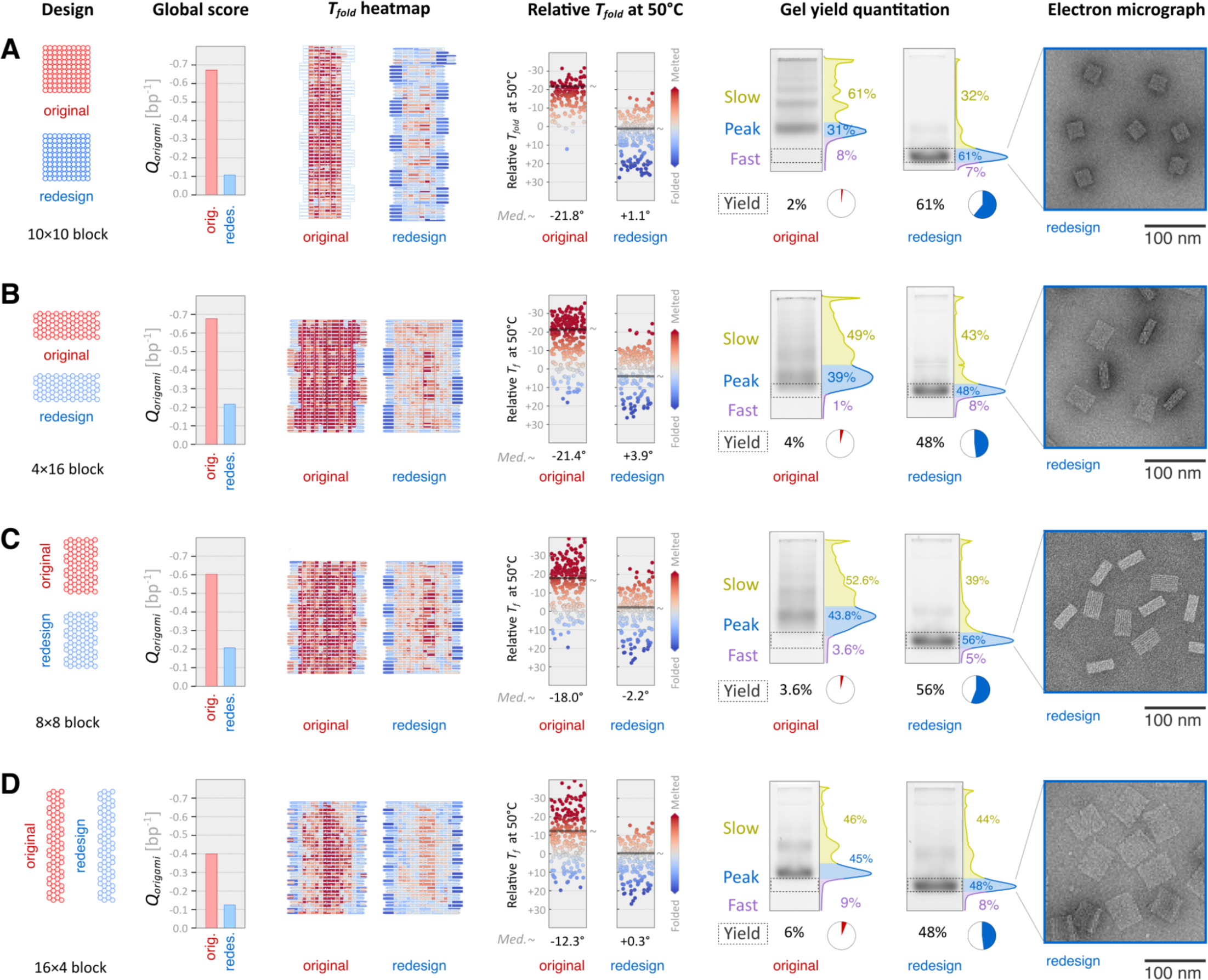
Experimental comparison of original and redesigned shapes. Four helix layouts were analyzed: (**A**) 10×10 square-lattice 100-helix block, (**B**) 4×16 block, (**C**) 8×8 block, and (**D**) 16×4 block. 64-helix blocks (B–D) used honeycomb lattice layouts. The first column shows the design cross-sections. The second column shows global quality scores per base pair (bp^-1^) (Eqn. 3). The third and fourth columns show two representations of per-staple folding temperature (*T_fold_*) estimates: a heatmap showing the staple routes and a strip plot with *T_fold_*values relative to 50°C, with ‘∼’ indicating median value. Staples predicted with *T_fold_* < 50°C are colored red, and *T_fold_* > 50°C are colored blue. The fifth column shows electrophoretic mobility analysis of samples folded in the presence of 20 mM MgCl_2_. ‘Yield’ is the integrated intensity of boxed regions divided by the total intensity of the lane. ‘Peak’ regions for redesigns were physically extracted and imaged by TEM; representative micrographs are shown in the sixth column.

Accurately folded DNA origami shapes tend to migrate on agarose gels as sharp, well-defined bands with increased mobility, while misfolded shapes migrate as diffuse bands with lower mobility along with additional species that indicate aggregation^5^. To assess folding accuracy and robustness, we folded each design at magnesium chloride (MgCl_2_) concentrations varying from 6 to 20 mM (Fig. S6), and quantified the 20 mM condition using ImageJ^28^.

We isolated a rectangular region of the lane spanning vertically from the bottom of the well down to the area below the fastest-moving band for each design pair, generated a 1D histogram, and manually subdivided the histogram into three regions: Slow, Peak, and Fast. Peaks correspond to the manually determined regions containing the leading band in each lane. Slow and Fast regions run above and below a Peak, respectively. We defined the Yield of accurate folding for each design as the integrated pixel intensity corresponding to the fastest moving Peak among the original and redesign conditions, which in all cases was the redesign. Our redesigned structures had uniformly better accurate-folding Yields compared to the originals. The 10×10 block redesign improved from 2% to 61% of band intensity qualified as accurate (Fig. 3A). The 64-helix blocks with 4×16, 8×8 and 16×4 configurations improved from 4% to 48%, 3.6% to 56%, and 6% to 48% (Fig. 3B–D). All redesigned variants displayed some degree of accurate folding at MgCl_2_ concentrations as low as 10 mM (Fig. S6).

We physically extracted Peak bands from the lanes containing the redesigned shapes and imaged them via TEM to confirm that the structures were intact and accurately folded (Fig. 3, column 5 and Figs. S7–S10).

### Key design principles

The original and redesigned shapes exhibited folding yields that correlated with global *Q* scores provided by our algorithm (Eqn. 3; Fig. 4A). To identify key design principles and related tunable design parameters, we analyzed the original and redesigned staple routes to identify the relative contributions of the *ΔG_hyb_*, *ΔG_loop_*, and *ΔG_bind_* to the overall *ΔG_total_*score.

**Figure 4.**
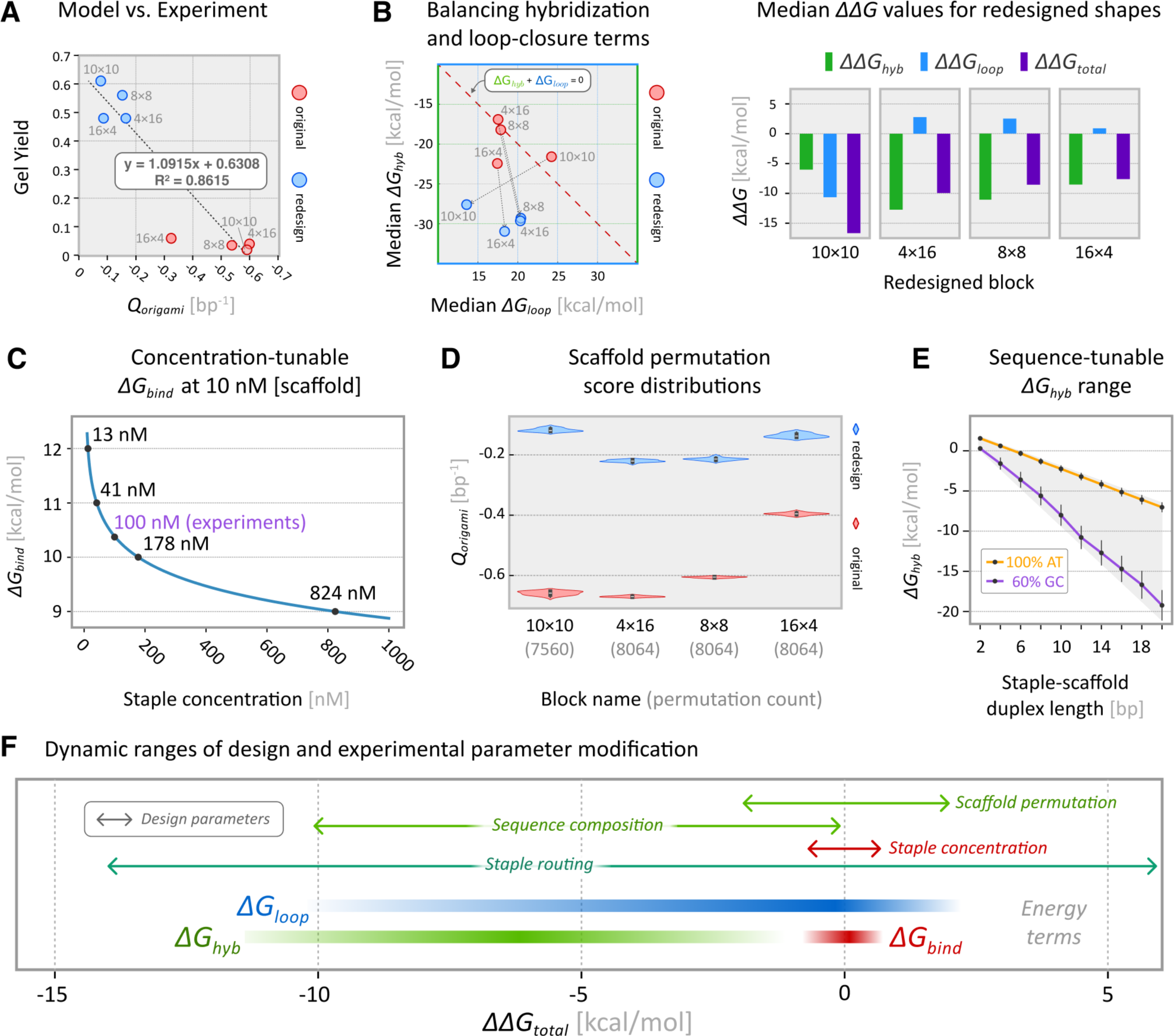
Analysis and modeling of key design parameters. (**A**) Global normalized design scores and gel yields correlate with R^2^=0.8615. (**B**) Plots of mean *ΔG_hyb_* and *ΔG_loop_*values for original and redesigned blocks show the relative contributions of hybridization and loop-closure to *ΔG_total_*. Constant *ΔG_bind_*values are not shown; staple concentrations were fixed across all folding reactions. Bar plot shows *ΔΔG* terms, or the change in median *ΔG* values between the original and redesigned shapes. (**C**) Increasing the concentration of a staple relative to the scaffold can marginally reduce its *ΔG_bind_* penalty term. (**D**) Global *Q_origami_*scores of every scaffold permutation form tight distributions. (**E**) Customizing the length and sequence of duplexes formed between staple segments and the scaffold provides a wide range of possible *ΔG_hyb_* values. (**F**) 1D plot of per-staple dynamic ranges, in change in kcal/mol, of user-designable parameters and corresponding energy terms from our thermodynamic model.

#### Balancing hybridization and loop-closure terms

Accurate folding requires staple routes that balance the enthalpically favorable formation of staple-scaffold duplexes via hybridization (*ΔG_hyb_*) with entropic penalties of scaffold loop-closures (*ΔG_loop_*). Fig. 4B shows the median *ΔG_hyb_* and *ΔG_loop_* values and how they changed between the original and redesigned shapes. Per-staple thermodynamic plots are shown in Fig. S11. Our model enables the quantitative comparison of specific design choices, including scaffold routing and passivation strategies. In the case of the 10×10 brick, both *ΔG_hyb_*and *ΔG_loop_*values improved markedly in our redesign (Fig. S2). For the remaining architectures, our algorithm determined staple routes with improved *ΔG_hyb_* values (*ΔΔG_hyb_* ranged from -8.5 to -12.7 kcal/mol), at the cost of slightly less favorable *ΔG_loop_* terms (*ΔΔG_loop_* of +0.9 to +2.8 kcal/mol).

#### Staple Concentration

The first term in our model, *ΔG_bind_*, accounts for the relative concentrations of scaffold and staple strands. For all lab tests we used fixed concentrations of 10 nM scaffold and 100 nM for each staple, therefore all edge weights included an identical *ΔG_bind_* value (10.3 kcal/mol). Our model predicts that further increasing a staple’s concentration, for example from 100 nM to 1000 nM will reduce its *ΔG_bind_* penalty by about 1.5 kcal/mol (Fig. 4C). Increasing the molar ratio of staples to scaffold above 10:1 is unlikely to significantly improve folding accuracy. In some cases, it may be worthwhile to boost individual staple concentrations, but the strategy is limited by practical considerations such as sample viscosity.

#### Scaffold Permutation

The sequence register of circular ssDNA scaffolds can be permuted within a design without modifying the staple routes. Staple *ΔG_hyb_* terms can vary across permutations, affecting the global quality score, *Q_origami_*. We modeled the effect of permuting the scaffold sequence to every possible register of each original and redesigned structure (Fig. 4D, Fig. S12A), and observed max-min spreads of up to 3.4 kcal/mol between the median *ΔG_hyb_* terms, and spreads up to 0.03 in *Q_origami_* score. We conclude that post staple-routing permutation analysis may improve folding accuracy, but the benefit is small relative to optimizing the staple routes. For comparison, the *Q_origami_*values in our redesigned shapes improved by 0.27 (16×4 block) to 0.56 (10×10 block) per base pair.

#### Scaffold Sequence

To assess the prospect of leveraging design-specific custom scaffold sequences to enhance folding accuracy, we modeled the sequence-tunable dynamic range of the *G_hyb_* term (Fig. 4E). We generated two sets of random duplex sequences with lengths varying from 4 to 20 nucleotides and GC content of 0–60%, the typical range offered by commercial DNA synthesis vendors. *G_hyb_* values of a 14-nt duplex, for example, differed by up to 8 kcal/mol based on GC content, suggesting that duplex-level sequence customization may offer a powerful complement to staple-breakpoint optimization for enhancing folding accuracy. Sequence customization will require design and synthesis of design-specific scaffolds.

Fig. 4F summarizes the potential *ΔΔG_total_*, in kcal/mol, that can be achieved by modifying key design and experimental parameters, along with *ΔΔG* ranges for the individual energy terms in our staple scoring function (Eqn. 4).

## Discussion

DNA origami design, like *de novo* protein design, presents a vast solution space. Unlike proteins, the geometric rules of DNA origami self-assembly have allowed for designing shapes without modeling the energetics of the folded state. However, the increasing demand for highly precise and repeatable fabrication in DNA origami applications signals an evolution toward computational modeling approaches analogous to those used in protein design.

Our study presents design principles that enhance the folding accuracy of DNA origami by integrating a global optimization algorithm with a robust thermodynamic model. The predictive capability of our scoring function, rooted in the thermodynamic stability of staple-scaffold interactions, has been empirically validated through electrophoretic mobility analysis and TEM imaging. In addition to the shapes reported here, our design principles have been successfully applied to generate precision instruments for probing cell activation and protein structure^29,30^.

Our methodology and computational toolset promise to broaden the horizons for the design of nanoscale constructs and pave the way for sophisticated applications previously challenging to achieve, from medical to electronic and photonic nanodevices. As an immediate practical benefit, improved folding accuracy and yield will lower costs for large-scale DNA origami manufacturing.

## Methods

### Origami design, optimization, and fabrication

Origami were designed in Cadnano^24^. Staple route auto-breaking, and analysis were performed using a custom software toolkit; Colab notebook available at the URL below. Folding reactions included 10 nM scaffold ssDNA (Tilibit) and 100 nM each staple (IDT). and 1XFOB20 (5 mM Tris-Base, 1 mM EDTA, 5 mM NaCl, 20 mM MgCl_2_ at pH 8.0). Magnesium-dependent stability was assessed using folding buffers containing 6 to 20 mM MgCl_2_ (Fig. S6). Folding reactions were performed using Bio-Rad MJ Research PTC-240 Tetrad thermal cycler. We used the following temperature annealing ramp: (1) Incubate at 65 °C for 10 min; (2) Incubate at 60 °C for 1 h, decrease by 1.0 °C every cycle; (3) Goto step 2 an additional 20 times.

### Gel analysis

Origami were analyzed using 2% agarose gel electrophoresis in Tris-borate-EDTA (45mM tris-borate and 1 mM EDTA) supplemented with 11 mM MgCl2 and SYBR Safe. Upon sample-loading, gels were run for 3 h at 80 V and subsequently scanned using a Typhoon FLA imager. Redesigned structures were purified by extracting the desired gel band using a razor blade, muddled to break down the agarose and then centrifuged through a Freeze ‘N Squeeze column (Bio-Rad).

### TEM characterization

For negative-stain grid preparation 5 μl of gel-purified DNA origami sample was deposited onto a glow-discharged thin-carbon coated grid, incubated for 1 min. Excess liquid was wicked away using filter paper. Next, 10 μl of freshly prepared 2% uranyl formate (Electron Microscopy Sciences) was applied to the grids and immediately wicked away using filter paper. A second round of 10 μl of 2% uranyl formate was then applied to the grids for 3 min before excess liquid was wicked away and the grid left to dry. Micrographs of the negatively stained grids were collected on Tecnai T12 (FEI) at ×30,000 magnification.

### Code Availability

The source code is available under an open-source license at: https://github.com/douglaslab/pyOrigamiBreak

A computational notebook is available at: https://colab.research.google.com/drive/1oavYcHN08N5lSMS1i5dtf-f3-j8dEe0-

## Acknowledgements

This work was supported by ONR grant N00014-17-1-2627, NSF grant DBI-1548297, and NIH grant R35GM125027. T.A. was supported by the Ruth L. Kirschstein NRSA Postdoctoral Fellowship grant F32GM119322.

## Author contributions

T.A. and S.M.D. conceptualized the project. T.A. performed research, collected data, wrote software and analyzed data. E.N. performed research and collected data. N.F. provided software support. S.M.D. provided resources, supervised the project, wrote software, analyzed data, and wrote the manuscript with input from all authors.

## Competing Interests

The authors declare no competing interests.

## Materials & Correspondence

Address correspondence to.

## Supplementary Information for

**Figure S1.**
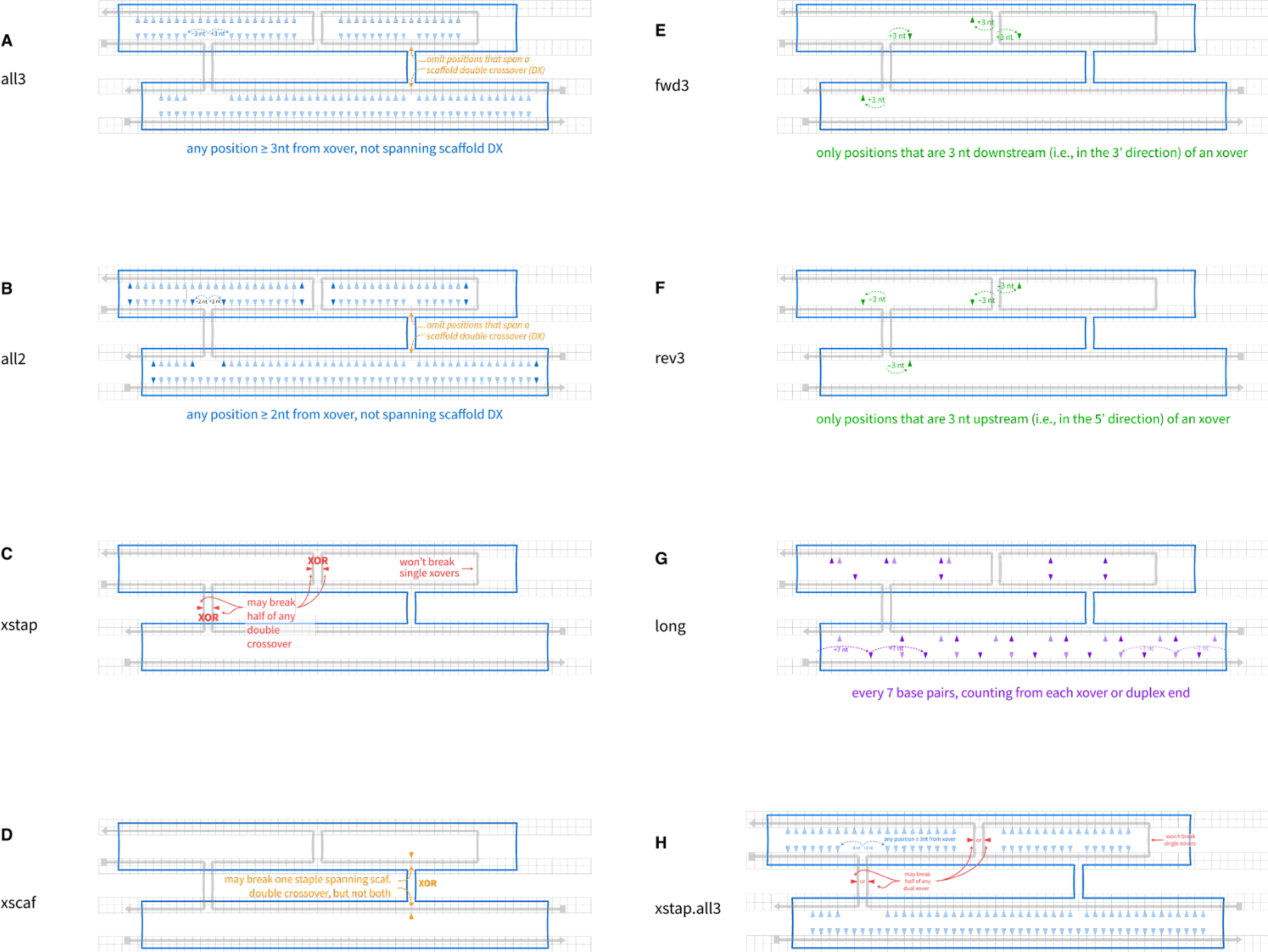
Breakpoint Rules. To identify the design principles driving folding accuracy, we needed a deterministic approach to encode and compare design rules. We devised a rule-based system for specifying the allowed staple ‘breakpoints,’ or locations where a precursor staple route is broken between two nucleotides, leaving a pair of 5’ and 3’ endpoints. Breakpoint rules are used in the construction of the weighted graph (Fig. 1A). We examined the following rules: (**A**) Rule ‘all3’ includes any breakpoint position ≥ 3nt from a staple crossover (xover), and not spanning a scaffold double crossover (DX). (**B**) Rule ‘all2’ includes any breakpoint position ≥ 2nt from a staple xover, and not spanning a scaffold DX. (**C**) Rule ‘xstap’ allows for breaking half of any staple DX. (**D**) Rule ‘xscaf’ allows for breaking one of the staples spanning a scaffold DX, but not both. (**E**) Rule ‘fwd3’ includes only positions that are 3 nt downstream (i.e., in the 3’ direction) of an xover. (**F**) Rule ‘rev3’ includes only positions that are 3 nt upstream (i.e., in the 5’ direction) of an xover. (**G**) Rule ‘long’ includes a breakpoint every 7 nucleotides pairs, counting from each xover or duplex end. (**H**) Rule ‘xstap.all3’ is a composite rule that includes the union of all breakpoints allowed by the ‘xstap’ and ‘all3’ rules.

**Figure S2.**
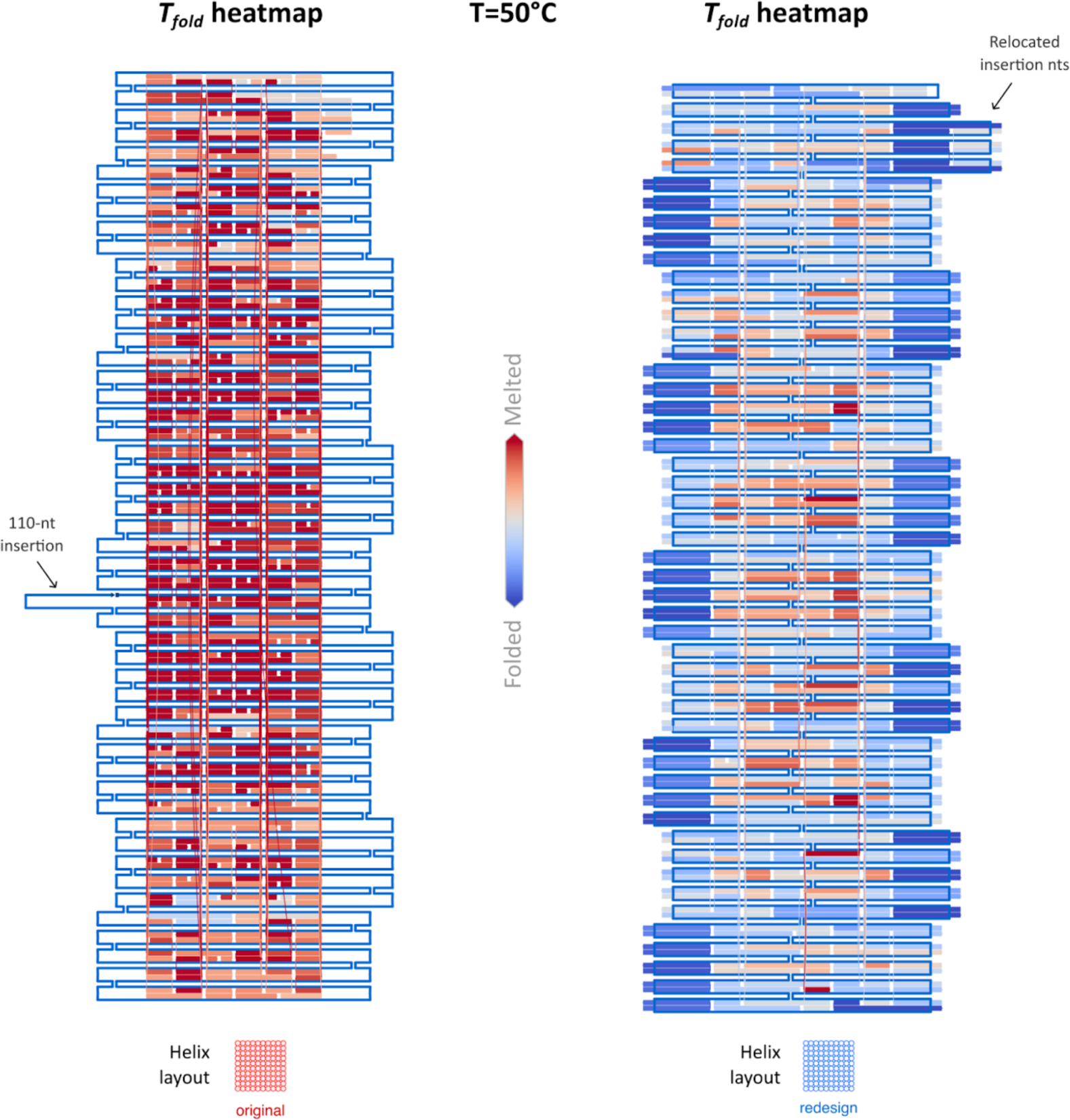
10×10 block heatmaps. To mitigate aggregation, the original design relied on a passivation strategy of leaving unpaired scaffold nucleotides at the left and right edges. A total of 2725 scaffold nucleotides are left unpaired, including a 110-nt ‘insertion’ at the helix[base] position 56[30]; the remaining 4835 scaffold bases form duplexes with staple strands. In our redesign, we rerouted the internal scaffold crossovers to a ‘middle-seam’ layout, and relocated the 110-nt insertion domain to extend helices 4, 5, 6, 7, 8, and 9 on the right edge. Before applying our staple-routing algorithm, we added edge staples that utilize unpaired ‘TTT’ overhangs for aggregation, resulting in 2719 added base pairs (6 of 7560 remain unpaired). The added edge staples in the redesigned 10×10 block have low *ΔG_loop_* penalties due to the revised scaffold crossover layout, resulting in the overall decrease of the median *ΔG_loop_* scores shown in Fig. 4B.

**Figure S3.**
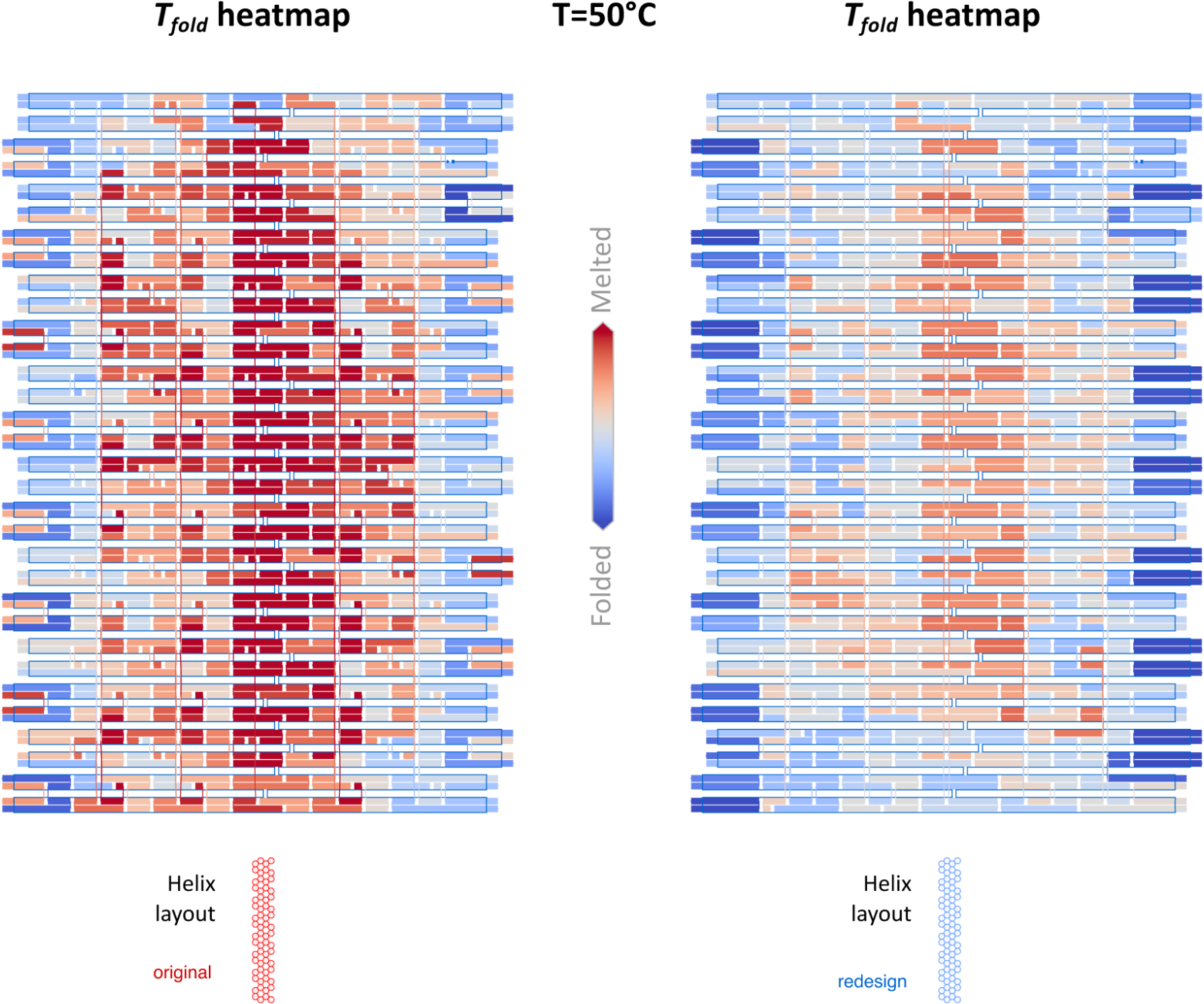
16×4 block heatmaps. The 16×4 blocks used identical scaffold routes and staple-passivation strategies.

**Figure S4.**
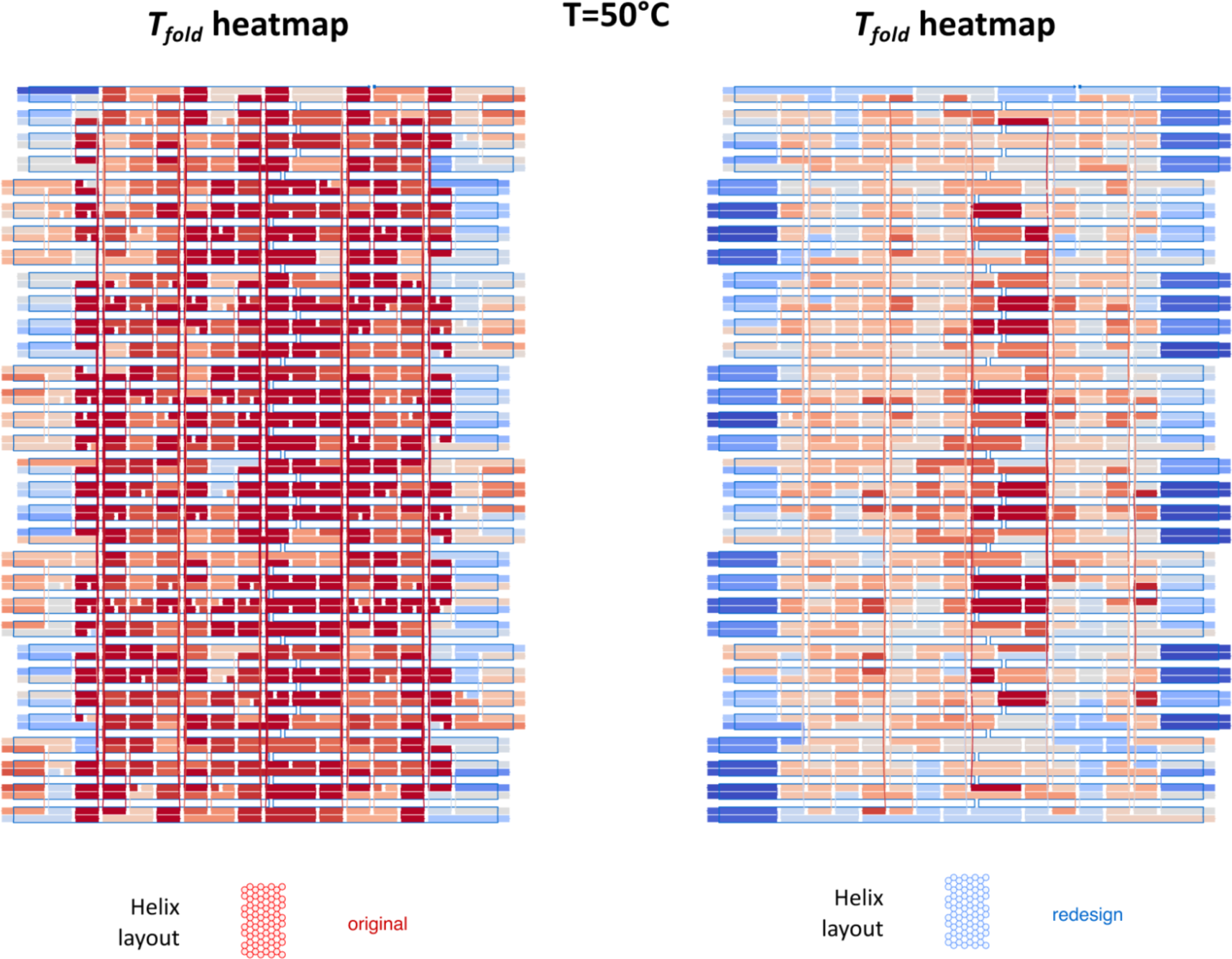
8×8 block heatmaps. The 8×8 blocks used identical scaffold routes and staple-passivation strategies.

**Figure S5.**
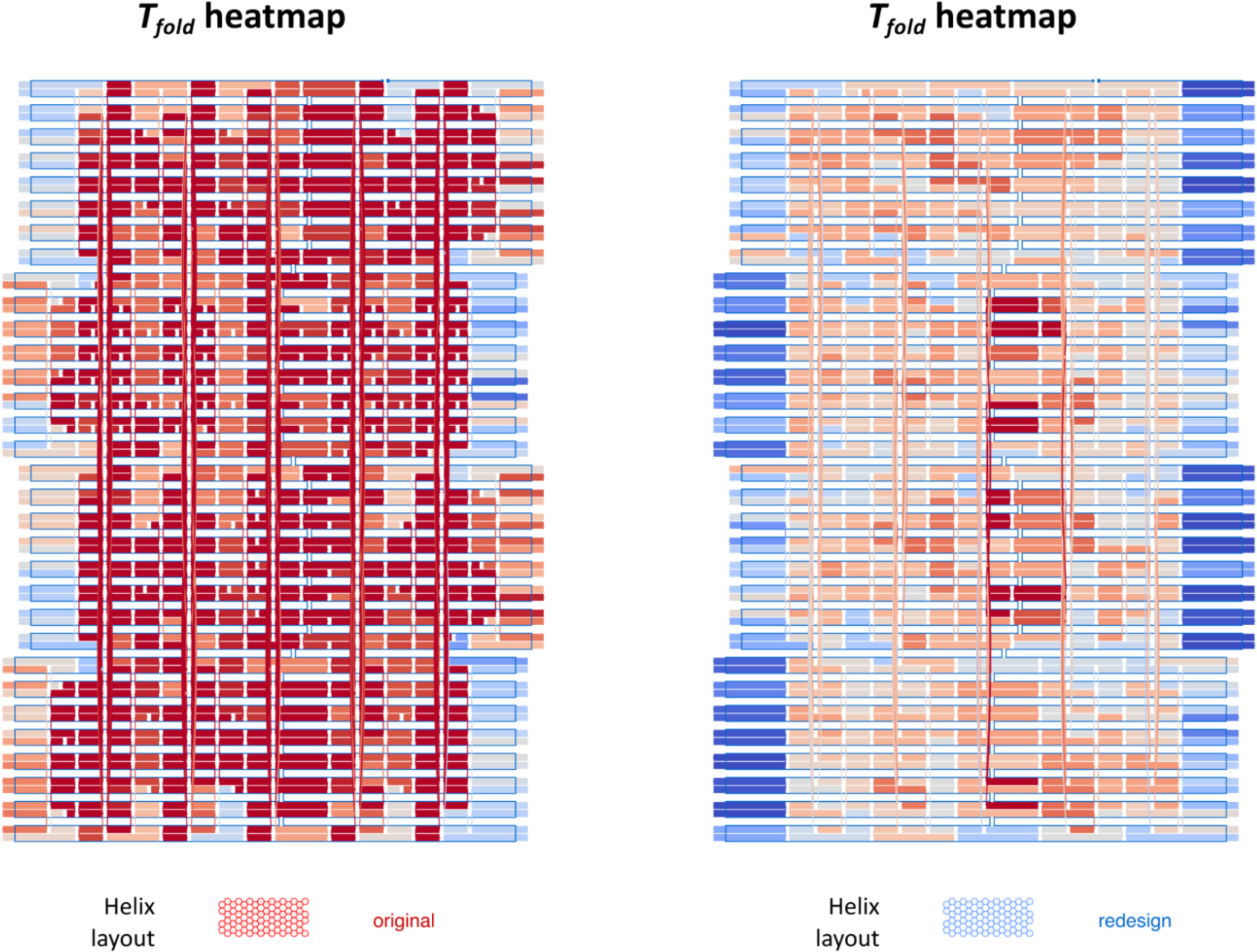
4×16 block heatmaps. The 4×16 blocks used identical scaffold routes and staple-passivation strategies.

**Figure S6.**
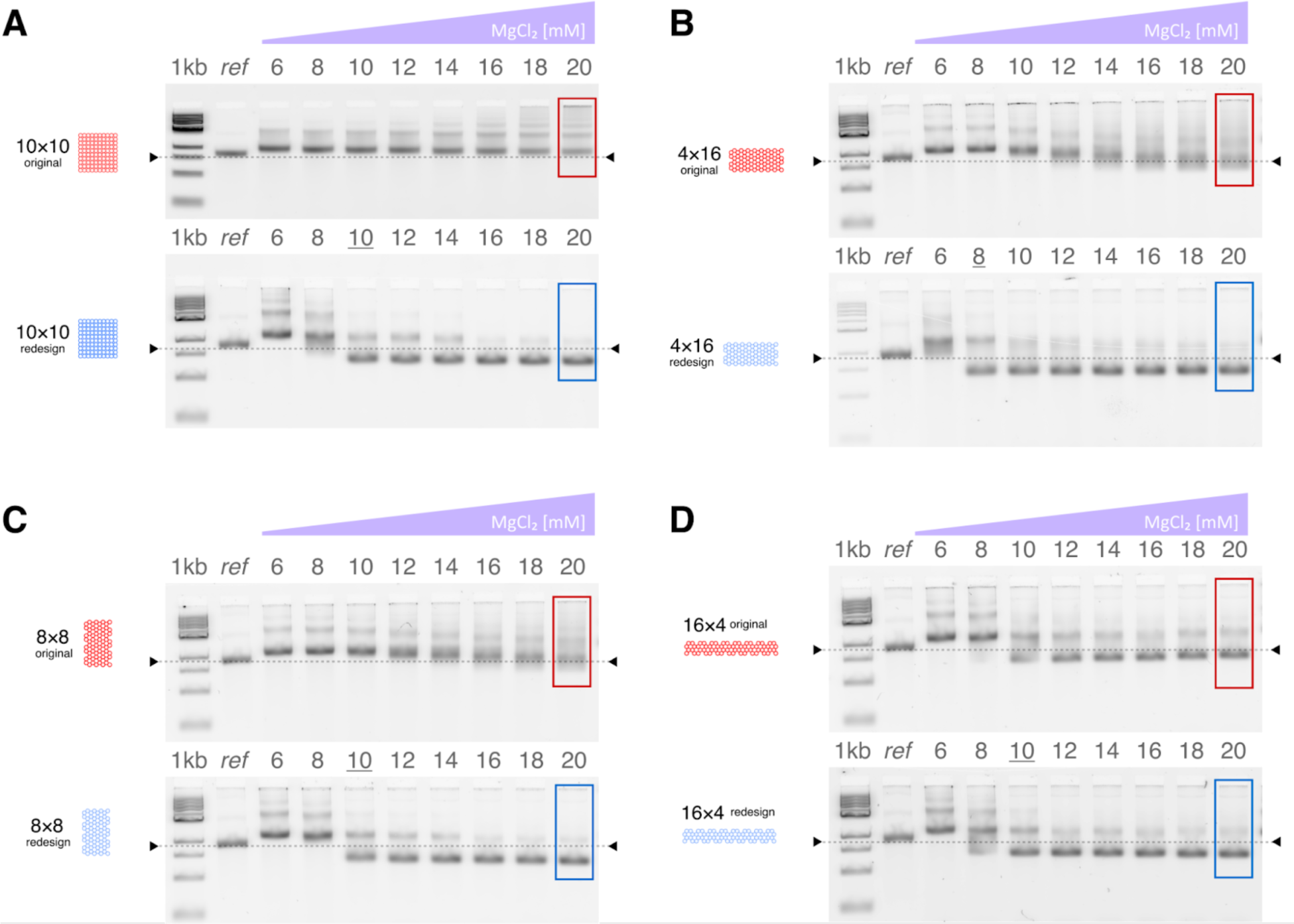
Detailed gel analysis. (**A**) 10×10 block. (**B**) 4×16 block. (**C**) 8×8 block. (**D**) 16×4 block. We folded each design and ran on separate gels as described in the Materials and Methods section. Lane 2 (“ref”) in each gel contained p7650 (A) or p8064 (B–D) scaffold ssDNA as a reference band for alignment and scaling during image analysis. Lanes 3–10 are labeled according to MgCl_2_ concentration. Intensity histograms of the boxed regions of the 20 mM MgCl_2_ lanes are shown in Fig. 3.

**Figure S7.**
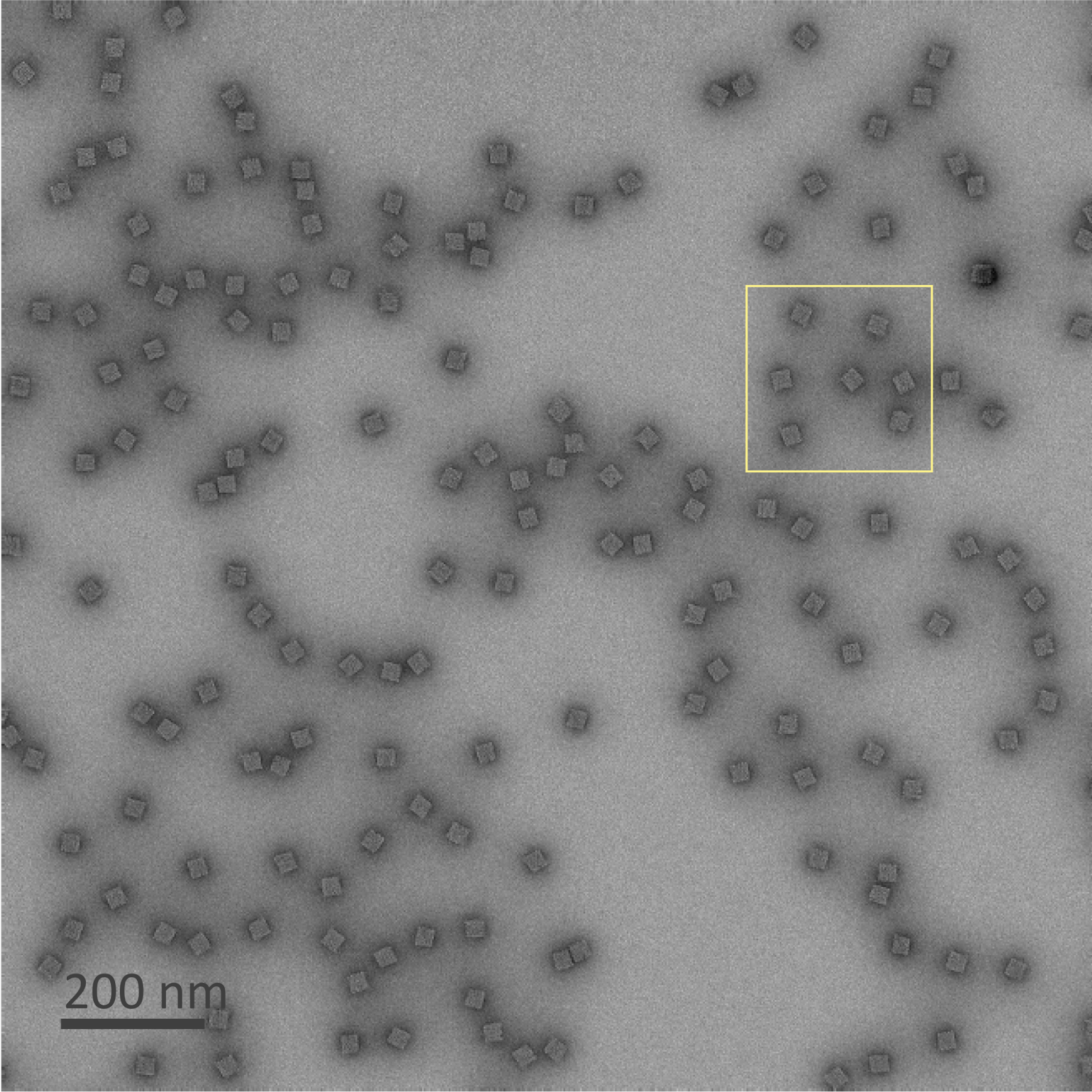
10×10 block TEM image. Negative stain. Boxed region is identical to Fig. 3A. Scale bar: 200 nm.

**Figure S8.**
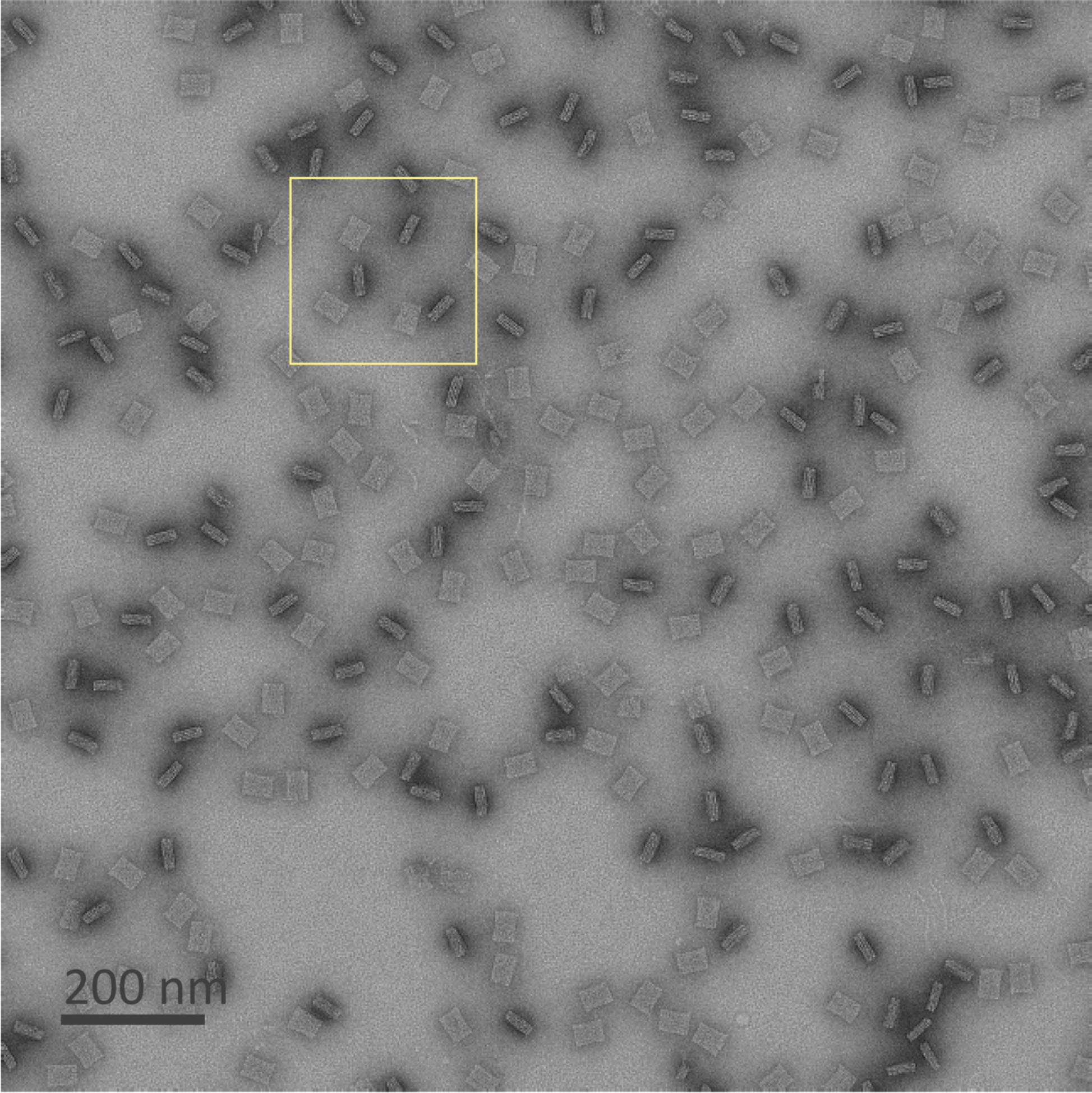
4×16 block TEM image. Negative stain. Boxed region is identical to Fig. 3B. Scale bar: 200 nm.

**Figure S9.**
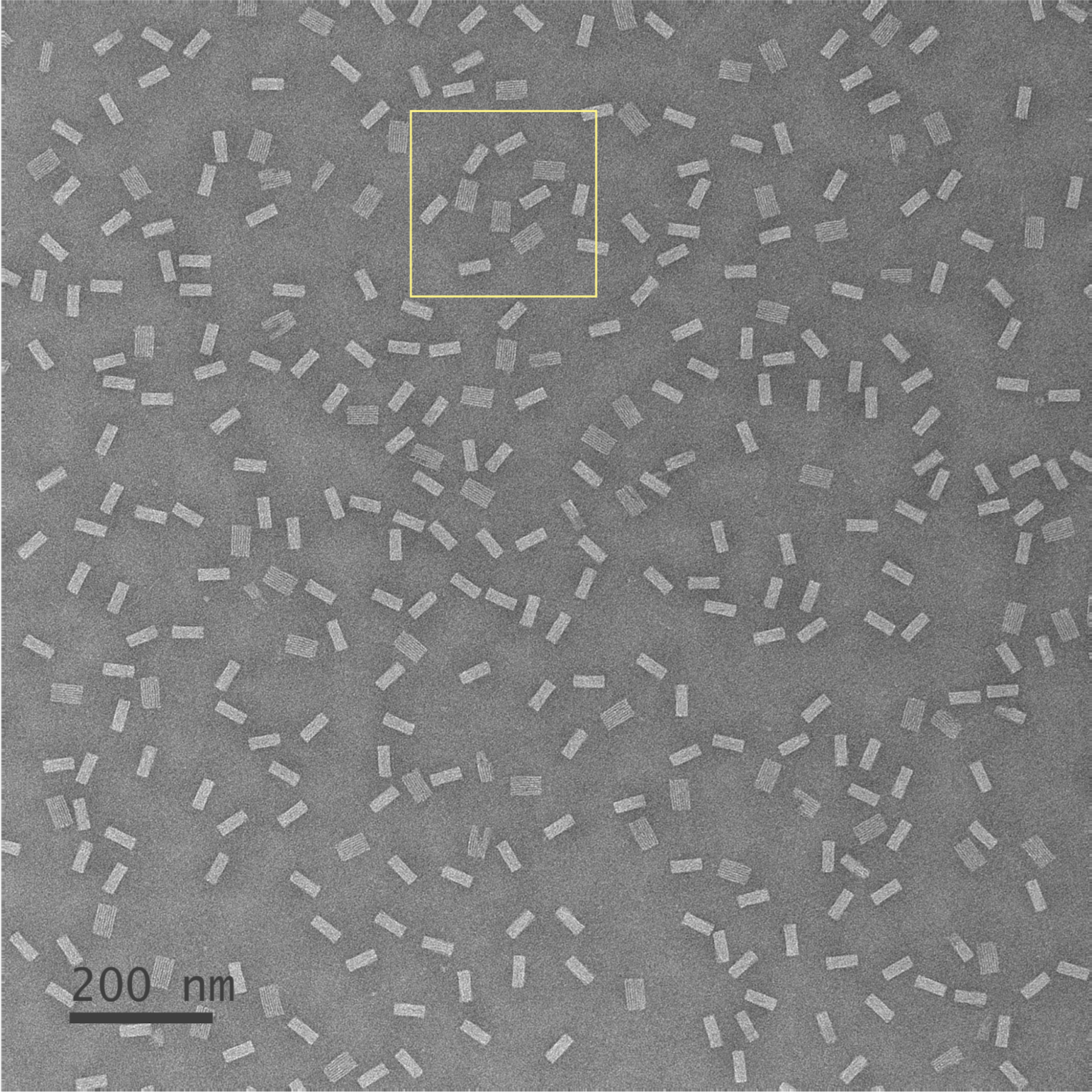
8×8 block TEM image. Negative stain. Boxed region is identical to Fig. 3C. Scale bar: 200 nm.

**Figure S10.**
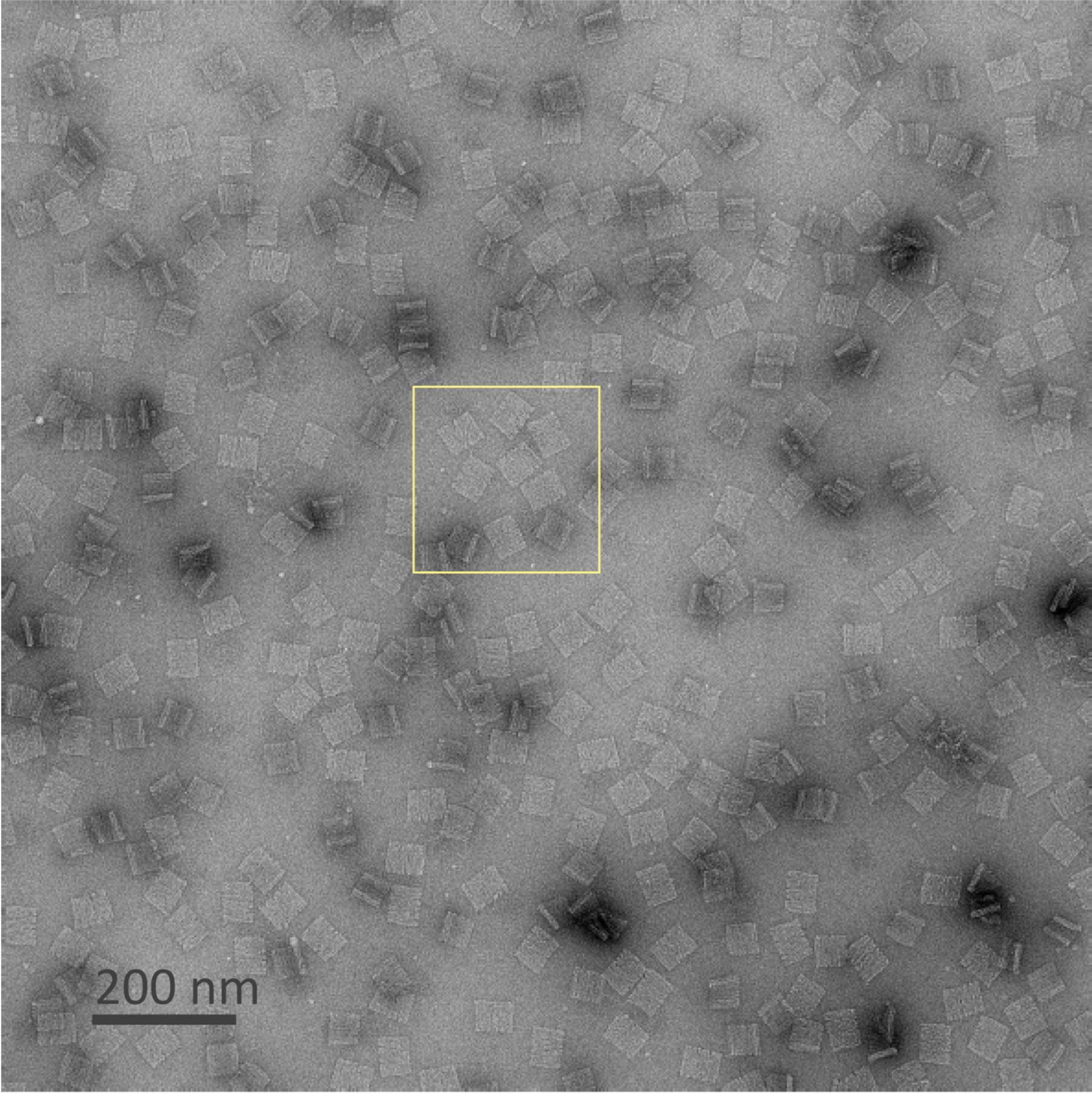
16×4 block TEM image. Negative stain. Boxed region is identical to Fig. 3D. Scale bar: 200 nm.

**Figure S11.**
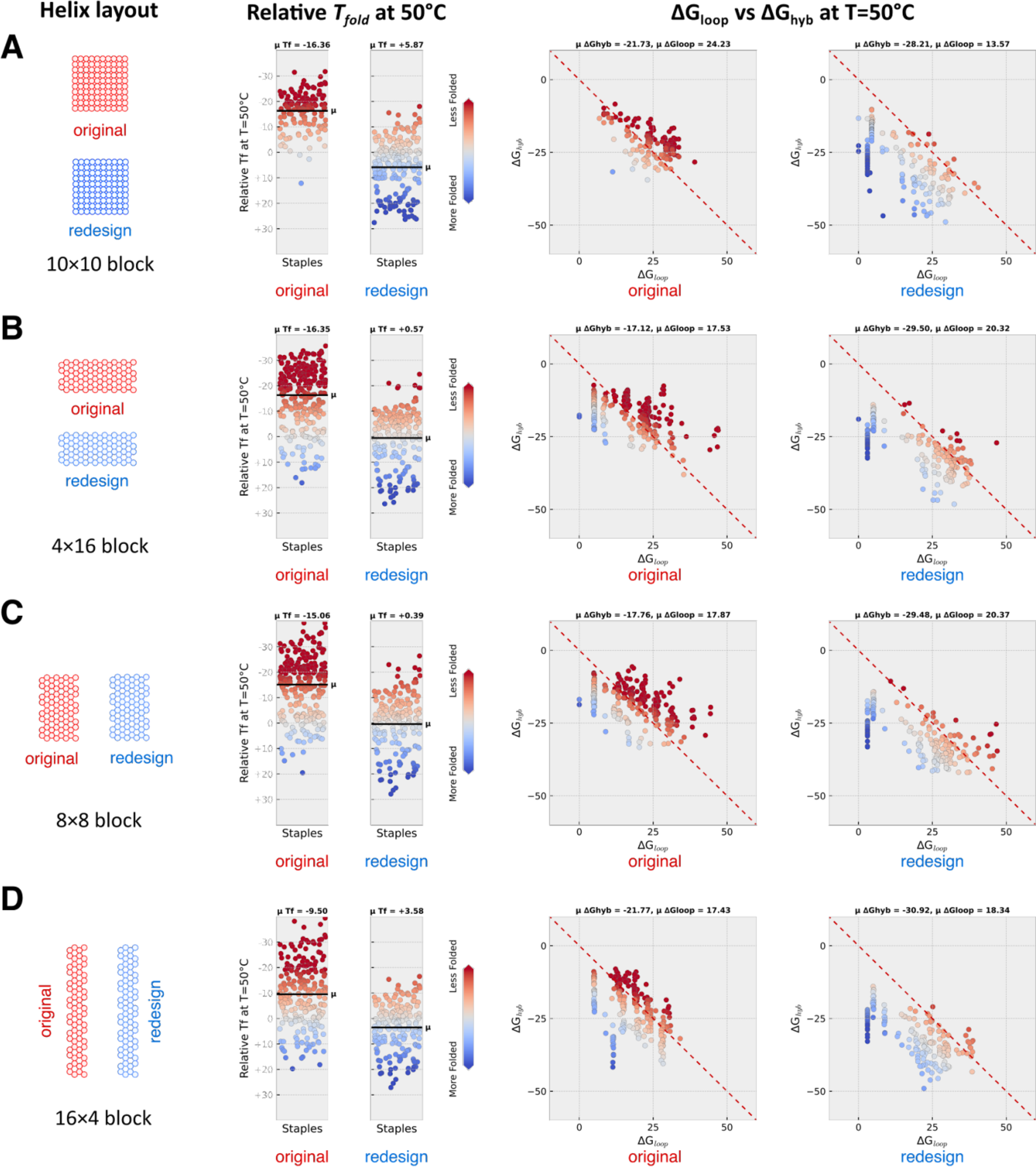
Per-staple thermodynamic analysis. (**A**) 10×10 block. (**B**) 4×16 block. (**C**) 8×8 block. (**D**) 16×4 block. Strip plots show relative *T_fold_*values at T=50°C. Color map ranges from +20°C (blue, more folded) to 0° (gray, 50% folded) to -20°C (red, less folded). Out-of-range values (>20°C or <-20°C) use the color of the closest in-range value. Mean (µ) *T_fold_* values are indicated by a horizontal black line. Scatter plots show per-staple *ΔG_loop_* (x-axis) and *ΔG_hyb_* (y-axis), with points colored according to their *T_fold_* values from the strip plots. Dashed line (red) indicates where *ΔG_loop_* + *ΔG_hyb_* = 0. Staples with values of *ΔG_loop_* + *ΔG_hyb_* < 0 appear below the diagonal and are predicted to fold at higher temperatures (blue points). The mid-range *T_fold_* points (gray) appear slightly off the diagonal due to the *ΔG_bind_* penalty term.

**Figure S12.**
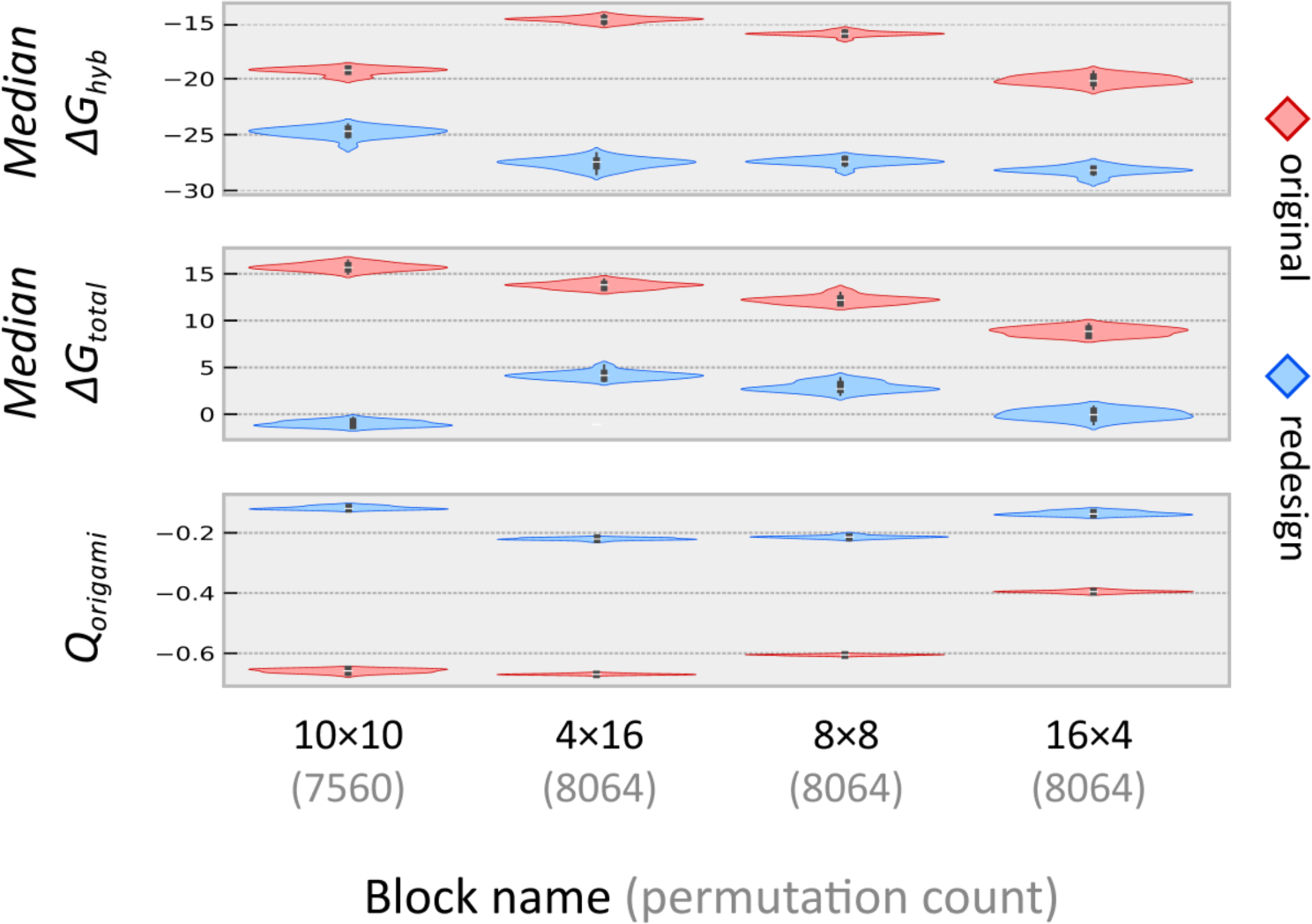
Scaffold permutation score distributions. We calculated *ΔG_hyb_* and *ΔG_total_* scores for each design variant across all circular scaffold permutations. The *nth* scaffold permutation is generated by removing the first *n-1* nucleotides of the sequence and appending them to the end of the sequence. The global *Q_origami_* values are reproduced from Fig. 4C for comparison.

## Notes

### Competing Interest Statement

The authors have declared no competing interest.

